# High-resolution mastigoneme structure reveals 5’,5’-phosphodiesters stabilized glycan folding

**DOI:** 10.1101/2024.12.24.630281

**Authors:** Junhao Huang, Hui Tao, Jikun Chen, Junmin Pan, Chuangye Yan, Nieng Yan

**Author notes:** To whom correspondence should be addressed: Nieng Yan; Chuangye Yan; Junmin Pan; Junhao Huang. These authors contributed equally (Junhao Huang, Hui Tao).

## Abstract

Glycans play a crucial role in structure organization, energy metabolism, and signal transduction in living organisms. Compared with proteins and nucleic acids, glycans exhibit remarkable molecular heterogeneity, extensive conformational flexibility, and diverse linkages. These properties pose significant challenges for obtaining high-resolution structure of glycans. Here, we report a cryo-EM structure of the heavily glycosylated *Chlamydomonas* mastigoneme at 2.3-2.5 Å resolutions. In addition to enabling analysis of accurate interactions for glycosyl packing, the high resolution map reveals an unprecedented 5’,5’-phosphodiester bond that links adjacent glycan chains attached to hydroxyproline (Hyp) residues n and n+3. Structural analysis reveals a secondary structural element for glycoconjugates, which we name the poly-Hyp (pHP) glycohelix. Our work represents an important advancement in deciphering glycan folding.

## Introduction

Carbohydrates, or glycans, are involved in a wide range of biological functions including metabolism, structural assembly, and signaling. Despite their fundamental significance, the three-dimensional structural information of glycans has been largely missing(1–5). To date, high-resolution structures of carbohydrates have been limited to protein-bound sugar ligands or short-chain oligosaccharides that are O- or N-linked to proteins(6–11). This lack of structural insights has significantly hindered our mechanistic understanding of the broad physiological and pathophysiological functions. Given that carbohydrates are the most abundant biomolecules on earth(1, 12, 13), an advanced comprehension of their higher-order glycan structures is essential for deciphering the folding principles of glycans and glycoconjugates, as well as shedding light on various research areas, such as materials sciences(14–20).

The challenge for obtaining high-resolution glycan structures stems from their diverse linkages, complex stereochemistry, and conformational heterogeneity(21–25). The remarkable flexibility of glycans hinders crystallization, rendering X-ray crystallography impractical for structural determination. Other conventional techniques, such as nuclear magnetic resonance, mass spectrometry, and scanning tunneling microscopy, are typically inadequate for resolving the structures of complex or low-abundance glycans(26–32). Moreover, the sample preparation process often destroys the native conformations of the glycans.

Over the past decade, single-particle cryo-electron microscopy (cryo-EM) has emerged as a transformative technique in structural biology(33–38). While cryo-EM has enabled the structural determination of many glycans conjugated to proteins, these studies have predominantly resolved glycans at moderate to low resolutions, often limited to regions proximal to the linked residues(39–44). One serendipitous exception came from the structural study of the non-tubular mastigoneme from *Chlamydomonas reinhardtii*(*45, 46*).

Mastigonemes are the lateral hairs that line the cilia of certain protists(45, 47–50), playing a crucial role in motility and mechanosensation(48–52). In the cryo-EM reconstruction of the stem region of the mastigoneme, approximately 25% of the molecular mass belongs to well-ordered glycans(45). In addition to the glycoprotein Mst1, which was the only known extracellular component of mastigonemes before the structure was resolved, the 3.0-Å resolution cryo-EM map revealed a large glycoprotein, which we named Mstax for Mastigoneme-specific axial protein (Fig. 1A).

**Fig. 1.**
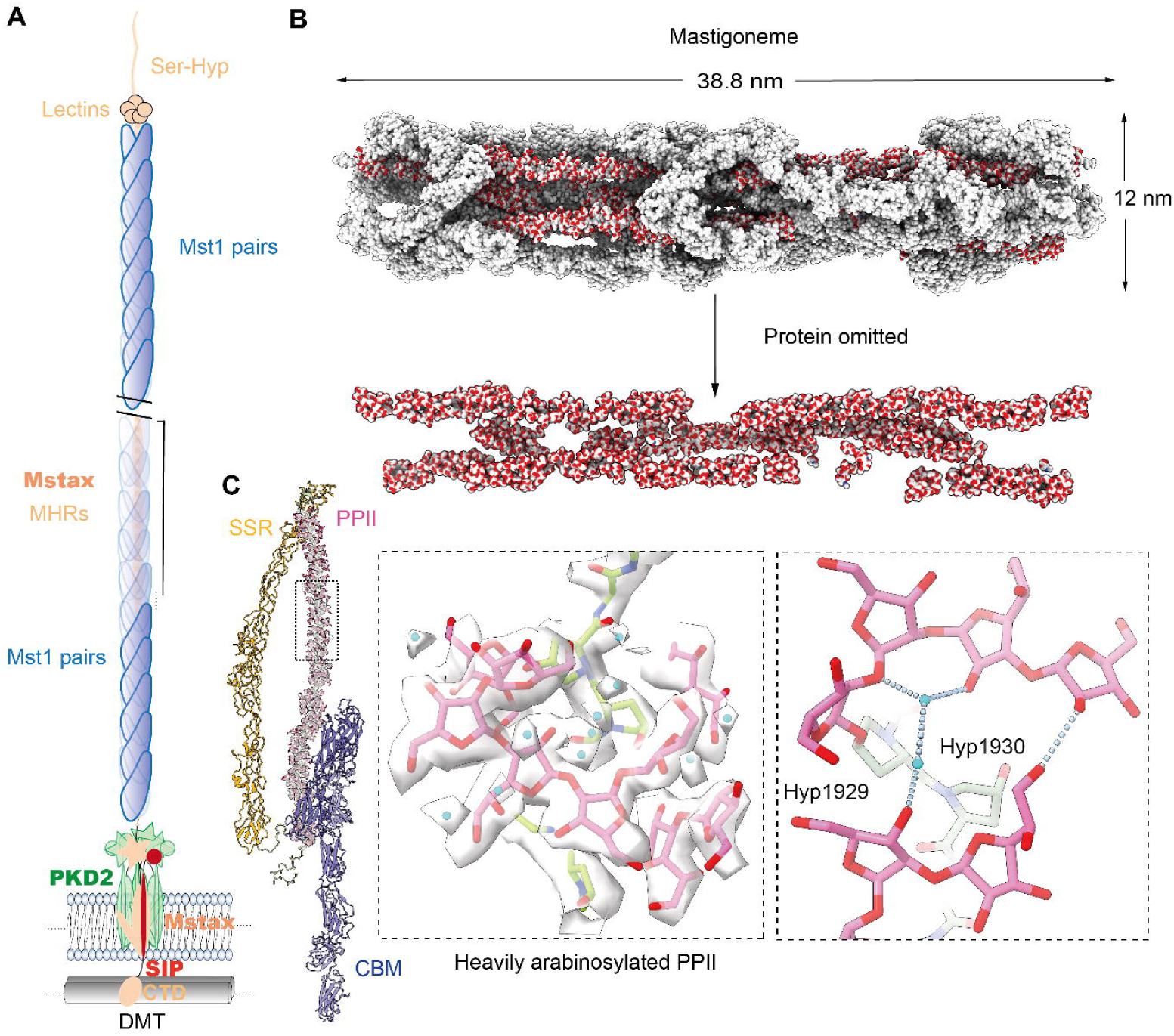
High resolution cryo-EM reconstruction of well-ordered glycans in native mastigonemes isolated from *Chlamydomonas reinhardtii*. **(A)** A schematic introduction to the overall assembly of the native mastigoneme in *Chlamydomonas reinhardtii*. A mastigoneme is constituted by one copy of Mstax that is surrounded by orderly assembled Mst1 proteins. The central Mstax, which consists of ∼8000 residues, spans the entire mastigoneme from the intracellular microtubule-interacting region to the distal tip. The predicted domains are labeled, including the Ser-Hydroxyproline (Ser-Hyp)-rich segment, 5 lectin domains, 34 Mstax-specific Hyp Repeats (MHRs), a PKD2-like transmembrane region that may interact with three PKD2 protomers and a single-span transmembrane protein SIP, and the cytosolic region that may associate with the microtubules. The stem of the mastigoneme is constituted by Mstax-MHRs and the surrounding Mst1 proteins that pack to super spirals. Please refer to Huang *et al* for the organization details of the mastigoneme(45). **(B)** Well-ordered glycans represent ∼ 25% molecular mass of the stem of a mastigoneme. Glycans on both Mstax and Mst1 are clearly resolved in the 3D cryo-EM map determined at resolutions of 2.26-2.48 Å. The overall structure, shown as spheres, of one segment of the mastigoneme stem that corresponds to two helical repeats is shown on the top. Protein moieties are colored silver, and the glycans are colored red for oxygen and silver for other elements. The protein-omitted structure is shown on the bottom to highlight the glycans. **(C)** Detail of the glycan chains in Mst1, including the bound water molecules, are revealed in the high-resolution map. Shown on the left is an updated overall structure of a Mst1 protomer. With 1987 residues in length, the signal peptide-cleaved Mst1 consists of 9 carbohydrate-binding modules (CBMs), 27 disulfide bond-containing repeats (SSR1-27), and a polyproline type II (PPII) helix (Ser-Hyp_n_ repeat rich). *Left inset*: EM map for a representative segment in the PPII helix. The PPII helix is constituted by Ser-Hyp repeats, wherein a typical 3-4 β-L-arabinofuranose (β-L-Ara*f*_3-4_) is O-linked to the hydroxyl group of each Hyp. Potential water molecules are shown as cyan spheres. *Right inset*: Representative water-mediated hydrogen bonds (H-bonds) in the PPII helix. Multiple hydrogen bonds are formed between sugar residues and nearby waters, both well resolved in the EM map. H-bonds are indicated with cyan, dashed lines. All EM maps and structure figures were prepared in ChimeraX(53).

Mstax, containing nearly 8000 residues, serves as the central shaft for the entire mastigoneme, with its amino (N) terminus located at the distal end. Following a Ser-hydroxyproline (Hyp)-rich segment and five lectin domains, residues 1408-3580 of Mstax form an extended linear polypeptide chain that is heavily glycosylated. This long thread, approximately 19.4 nm in length, contains 34 repeats of a unique sequence, Hyp_11_-(X_2_Hyp)_5-6_-Hyp3(X_2_Hyp)_2_-Hyp_12_ (short as Hyp_11_(X_2_Hyp)_n_ Hyp_12_), which we named the Mstax-specific Hyp Repeat (MHR). The MHR region, spanning roughly 660 nm, defines the stem of a mastigoneme (Fig. 1A).

Each Hyp residue is modified with well-ordered arabinoglycans, which play a major role in mediating the assembly of Mst1 around Mstax-MHR at a stoichiometric ratio of 2:1. These findings position the mastigonemes as a potential model system for advancing our understanding of glycan folding and structure. However, despite the decent resolution of 3.0 Å in our previous study, ambiguities remained in the assignment of sugar moieties(45), and precise interpretation of inter- and intramolecular interactions was hindered by a lack of details on the bound water molecules. To address these questions, we sought to improve the resolution of the *Chlamydomonas* mastigoneme structure.

Here we report the high-resolution structure of the mastigoneme, wherein the sugar moieties and nearby water molecules can be unambiguously assigned. Unexpectedly, the ∼2.3 Å map reveals numerous 5’,5’-phosphodiester bonds that connect neighboring arabinoglycans from the n and n+3 Hyp residues in Mstax-MHR. The 5’,5’-phosphodiester sugar linkages, previously unreported, could only be revealed at the high resolution. These findings not only afford unprecedented insight into glycan structures, but will also contribute to the rational design of biomaterials.

## Results

### High resolution cryo-EM structure determination of mastigonemes

To achieve higher resolution for the well-ordered native glycans in the mastigoneme of *Chlamydomonas reinhardtii*, we increased the concentration of the mastigonemes and prepared cryo-EM samples following our published protocol(45). We manually selected 3,847 out of 7,193 collected micrographs for the cryo-EM analysis of the mastigoneme fibril. Finally, cryo-EM maps at resolutions at 2.26-2.48 Å were obtained for most regions of the mastigoneme stem. The improved resolution enabled the unambiguous assignment of nearly all sugar residues, as well as a large amount of water molecules and cations. In each Mst1 molecule, 1907 residues and 233 sugar residues were assigned, while in each Mstax-MHR, 64 residues and 114 sugar residues were modeled. The overall structure, particularly for the protein moieties, remains the same as reported previously. Hereafter we will focus on the unprecedentedly high-resolution glycan structures for illustration (Fig. 1B).

Unlike the MHRs of Mstax, the majority of which is covered with thick glycans, the glycosylation of Mst1 mainly occurs on its elongated type II polyproline (PPII) helix, which is enriched in Ser-Hyp_n_ repeats, where n ranges from 1 to 9, typically Ser-Hyp_3_ (Fig. 1B, C). The updated cryo-EM map confirms our previous assignment of L-arabinofuranosyl (L-Ara*f*) and D-galactofuranosyl (D-Gal*f*) to these chunks of glycan densities. Notably, many interacting water molecules are observed mediating the interactions between sugar chains (Fig. 1C). For instance, it is now clearly observed that the 2’-OH groups of Ara*f*_1_ and Ara*f*_3_/Gal*f*_3_ interact with each other via water-mediated hydrogen bonds (H-bonds). Additionally, the sugar chains from Hyp residues at positions n and n+3 also interact through both direct and water-mediated H-bonds. These interactions occur throughout the entire PPII helix and contribute to the structural stability.

### Lateral linkage of the glycans through 5’,5’-phosphodiester bonds

A totally unexpected discovery from the high-resolution map concerns the glycans on the MHRs of Mstax. In our previous study, we primarily assigned Ara*f*_2_-Gal*f* to each Hyp in the unique pattern of Hyp_11_-(X_2_Hyp)_n_-Hyp_12_ as the core structure for each glycan chain(45). Consequently, the two consecutive poly-Hyp segments are coated with glycoshells, while the (X_2_Hyp)_n_ segment features glycan “brushes” lining one side (Fig. 2A). When the resolution was improved from 3.0 Å to 2.26 Å, it turns out that the densities for any two adjacent glycan branches on the MHRs are connected on the outer edge in a highly conserved pattern (Fig. 2A).

**Fig. 2.**
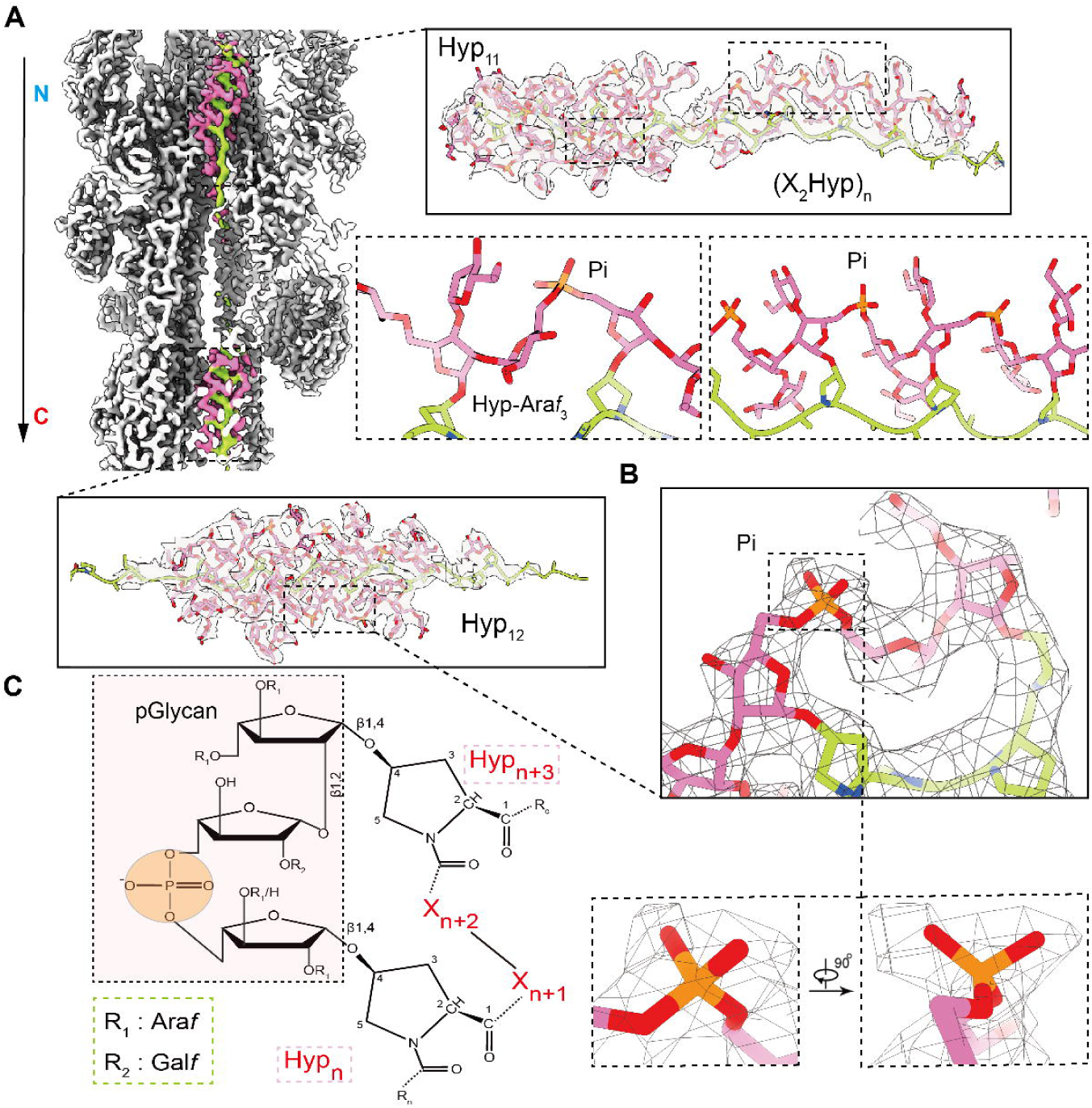
Stabilization of the glycan arrays in Mstax through unprecedented 5’,5’-phosphodiester bonds. **(A)** High-resolution glycan densities in Mstax reveal lateral arabinose stitching through 5’,5’-phosphodiester bonds. Shown on the left is the overall EM map of a segment that corresponds to one MHR in Mstax with surrounding Mst1 proteins (white). Each MHR comprises a heavily glycosylated Hyp_11_(X_2_Hyp)_n_ Hyp_12_ fragment that exists as an extended, linear peptide. Neighboring glycans that are respectively linked to Hyp residues n and n+3 in Hyp_11_-(X_2_Hyp)_n_ and Hyp_12_, with each glycan having a Ara*f*_2_-Gal*f* core, are connected by a 5’,5’-phosphodiester bond. In all panels, arabinoglycans and the MHR peptide are colored pink and green, respectively, and the phosphate is highlighted orange. **(B)** A typical density for the phosphate group. **(C)** Diagram of the chemical details of L-Ara*f* lateral linkage through 5’,5’-phosphodiester bond. The two L-Ara*f* moieties on the nearby Hyp residues n and n+3 are bonded to the phosphate group. R1 and R2 stand for Ara*f* and Gal*f*, respectively. X can be any residue, including Hyp as in the poly-Hyp segment. For description simplicity, we will describe the 5’,5’-phosphodiester bond-stitched glycans as the pGlycan.

The density for the linkage displays a tetrahedral contour that is characteristic of a phosphate group. When we modeled phosphate into this density, it was immediately evident that the neighboring arabinoglycans are linked through a phosphodiester bond, resembling the backbone linkages in DNA or RNA (Fig. 2B). This observation is further substantiated by the nuclear magnetic resonance (NMR).

Unlike the 3’,5’-phosphodiester bond in nucleic acids, the phosphodiester bonds in Hyp_11_-(X_2_Hyp)_n_ and Hyp_12_ are exclusively formed by the 5’-OH groups of two L-Ara*f* residues. Therefore, they are 5’,5’-phosphodiester bonds, which have never been reported in native glycans. This bond formation follows a consistent pattern, where the first L-Ara*f* (L-Ara*f*_1_) on Hyp_n_ is always connected to the second L-Ara*f* (L-Ara*f*_2_) on Hyp_n+3_ through the 5’,5’-phosphodiester bond (Fig. 2C). In addition, the third sugar, D-Gal*f*_3_ in L-Ara*f*_2_-D-Gal*f* is further confirmed by the high-resolution map. The glycan core can be further extended by adding a few Ara*f* residues on L-Ara*f*_1_ via its 2’-OH and 3’-OH groups (Fig. 2C).

Each Hyp residue in Hyp_11_ and Hyp_12_ shares the same glycosylation pattern, leading to three strips of 5’,5’-phosphodiester bonds lining the exterior of their glycoshells. Interestingly, the glycans attached to the Hyp residues in (X_2_Hyp)_n_ also follow the same pattern for phosphodiester formation. Since the first Hyp in (X_2_Hyp)_n_ is located third place from the last Hyp in Hyp_11_, their glycans are also connected by a 5’,5’-phosphodiester bond, in accordance with the abovementioned rule. In this way, the 5’,5’-phosphodiester bonds stitch together the neighboring glycans on the same side of the central peptide, forming a covalently bonded network. For description simplicity, we will refer to the 5’,5’-phosphodiester bond-linked glycans as the pGlycan (Fig. 2C).

### The pGlycans directly interact with Mst1

Our previous 3.0-Å resolution structure already suggested the important role of the glycans in mediating the assembly of Mst1 around Mstax-MHRs. The high-resolution maps reveal 5’,5’-phosphodiester bonds provide the molecular basis for the stable interactions between these glycans and the protein residues. Furthermore, the phosphate groups directly participate in binding with Mst1. We will not repeat our earlier findings(45), but focus on several phosphate-mediated interactions hereafter.

As described previously, we artificially divided the resolved mastigoneme repeat unit into three parts in the form of Mst1 pairs: pair 0, pair −1 and pair +1 (Fig. 3A). The Hyp_11_-(X_2_Hyp)_n_ segment mainly interacts with the disulfide bond-containing repeats (SSRs) 1-8 in Mst1 pair 0 and SSRs18-27 in pair +1.

**Fig. 3.**
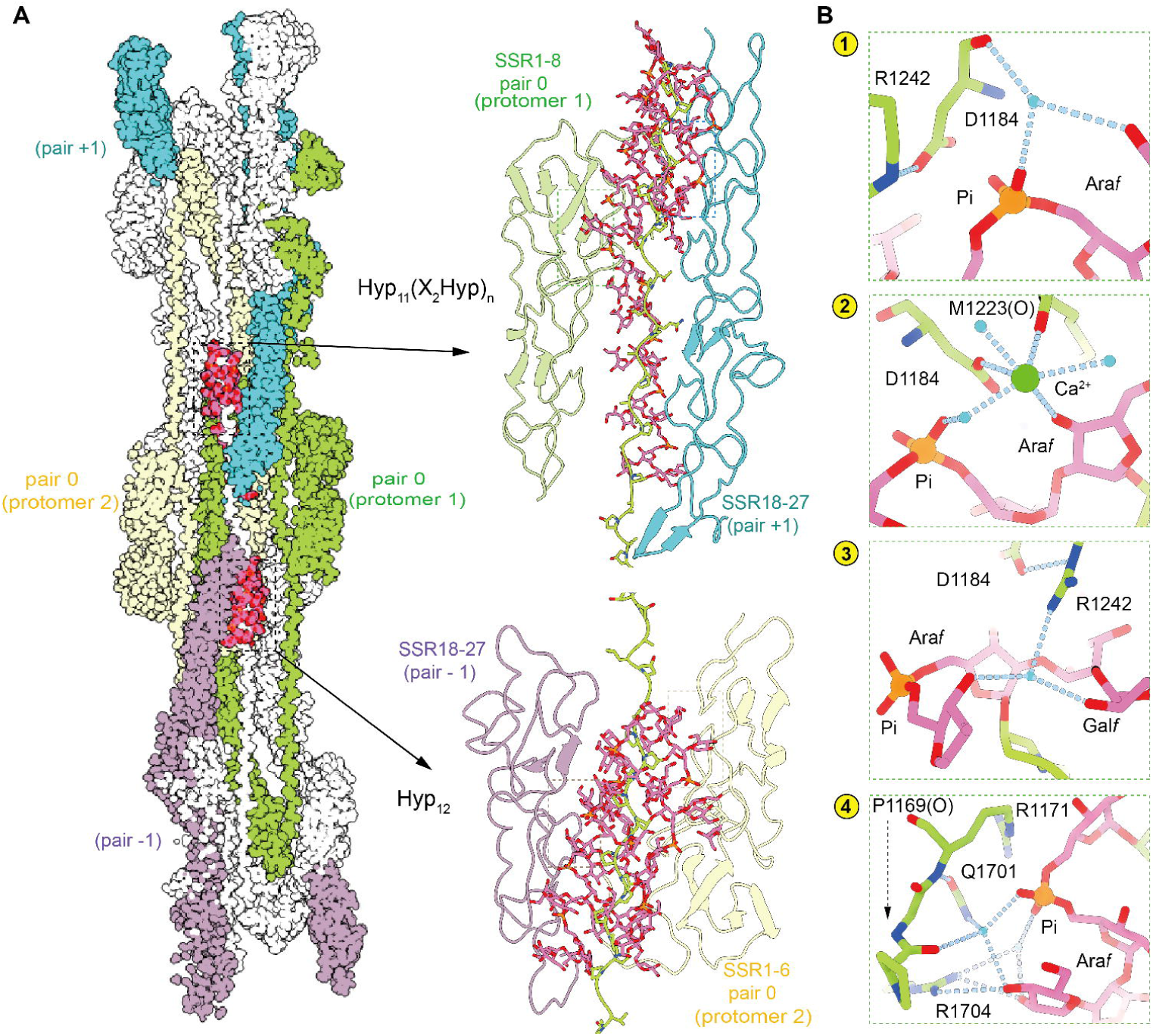
The 5’,5’-phosphodiester bond stabilized glycans mediate the assembly between Mstax and Mst1. **(A)** Hyp-linked glycans of Mstax engage in the interactions with surrounding Mst1 molecules. Shown here is the surface view of one helical repeat of a mastigoneme. Three pairs of Mst1 (−1, 0, +1) wrap around the MHR region of Mstax. Glycans on Hyp_11_-(X_2_Hyp)_n_ and Hyp_12_ exhibit nearly identical structures, and interact with the SSR domains from different Mst1 protomers in a similar manner. **(B)** Glycans medicate the assembly between Mstax and Mst1 through extensive polar interactions. Glycans on the Hyp_11_ and (X_2_Hyp)_n_ of Mstax interact with residues in SSRs 1-8 in Pair 0. The high resolution EM map allows for detailed analysis of their interactions. The phosphate group in the 5’,5’-phosphodiester linkage between L-Ara*f*s forms two sets of water-mediated H-bonds (insets 1 & 2). One forms an interaction with a potential calcium ion (green sphere). The SSRs 1-8 in Pair 0 on the other side also forms extensive interactions with the arabinose residue and phosphate group attached to Hyp_12_ (insets 3 & 4). Phosphorus atoms are colored orange.

The phosphate group in the 5’,5’-phosphodiester linkage between the L-Ara*f*_1_ on Hyp_11_ and the L-Ara*f*_2_ of the first (X_2_Hyp)_1_ on (X_2_Hyp)_n_ (short as Hyp_11_-(X_2_Hyp)_1_) forms two sets of water-mediated H-bonds. One set interacts with the carbonyl group of Asp1184 and the 5’-OH group of the branched L-Ara*f* on 3’-OH of L-Ara*f*_1_ from residue Hyp_11_. The other involves a water-mediated interaction with a cation that judged from the surrounding chemical groups, is likely to be calcium (Fig. 3B, insets 1 & 2). The cation is coordinated by the carboxyl group of Asp1184, the 3’-OH of L-Ara*f*_1_ of (X_2_Hyp)_1_, the carbonyl group of Met1223, and three water molecules. The guanidine group of Arg1242 stabilizes the carboxyl group of Asp1184 and forms a water-mediated interaction with both the 2’-OH of D-Gal*f*_3_ in (X_2_Hyp)_1_ and 3’-OH of L-Ara*f*_2_ in (X_2_Hyp)_2_ to help maintain the local conformation (Fig. 3B, inset 3).

The phosphate group in Hyp_7_-Hyp_10_ on the segment Hyp_11_ also establishes an extensive H-bond network with residues both from SSRs1-8 of pair 0 and SSRs18-27 in pair +1 (Fig. 3B, inset 4). This phosphate group is stabilized by the positively charged guanidium group of Arg1171. It also forms two water-mediated interactions. One water molecule mediates the interaction with the carbonyl group of Pro1169, the 3’-OH group of the branched L-Ara*f* on 3’-OH of L-Ara*f*_1_ from residue Hyp10, and the side chain of Gln1701. The other water mediates the interaction with the 2’-OH group of the same branched L-Ara*f* and Arg1704. Arg1704 also contribute to stabilizing this branched L-Ara*f*.

Hyp_12_ also interacts with the Mst1 molecule in pair 0 and pair −1 on the corresponding regions of the SSR, which matches the pseudo C2 symmetry assembly model of the mastigoneme. Interactions between the phosphate and glycan of Hyp_12_ are similar to those observed for Hyp_11_ (Fig. 3B).

### Common features of the glycan helix lining poly-Hyp (pHP-glycohelix)

In the high-resolution structure of the *Chlamydomonas* mastigoneme, there are three major clusters of arabinoglycans. One coats the PPII helix in Mst1 and lacks any phosphodiester bond, while the other two pGlycans on Mstax-MHR are in essence the same, one lining the (X_2_Hyp)_n_ in a single array, and the other surrounding Hyp_11/12_ as three arrays (Fig. 4A). A detailed analysis reveals that, regardless of the presence or absence of phosphodiester bonds, all these three clusters exhibit similar structural parameters when comparing the glycans attached to Hyp residues n and n+3. The glycans coating the PPII helix in Mst1 can be superimposed with those surrounding Hyp_11/12_, which are literally three parallel strands of those lining (X_2_Hyp_2_)_n_ (Fig. 4B). In this sense, the glycans on Hyp residues n and n+3 constitute one secondary structural element. Each helical turn spans ∼3.2 glyco-Hyp with an average twist and rise of ∼112.5° and 3.2 Å, respectively, from Hyp residues n to n+1. Thus, in one array of the glycan chain, the average twist and rise will be ∼22.5° and 9.6 Å, respectively, from Hyp residues n to n+3 (Fig. 4A).

**Fig. 4.**
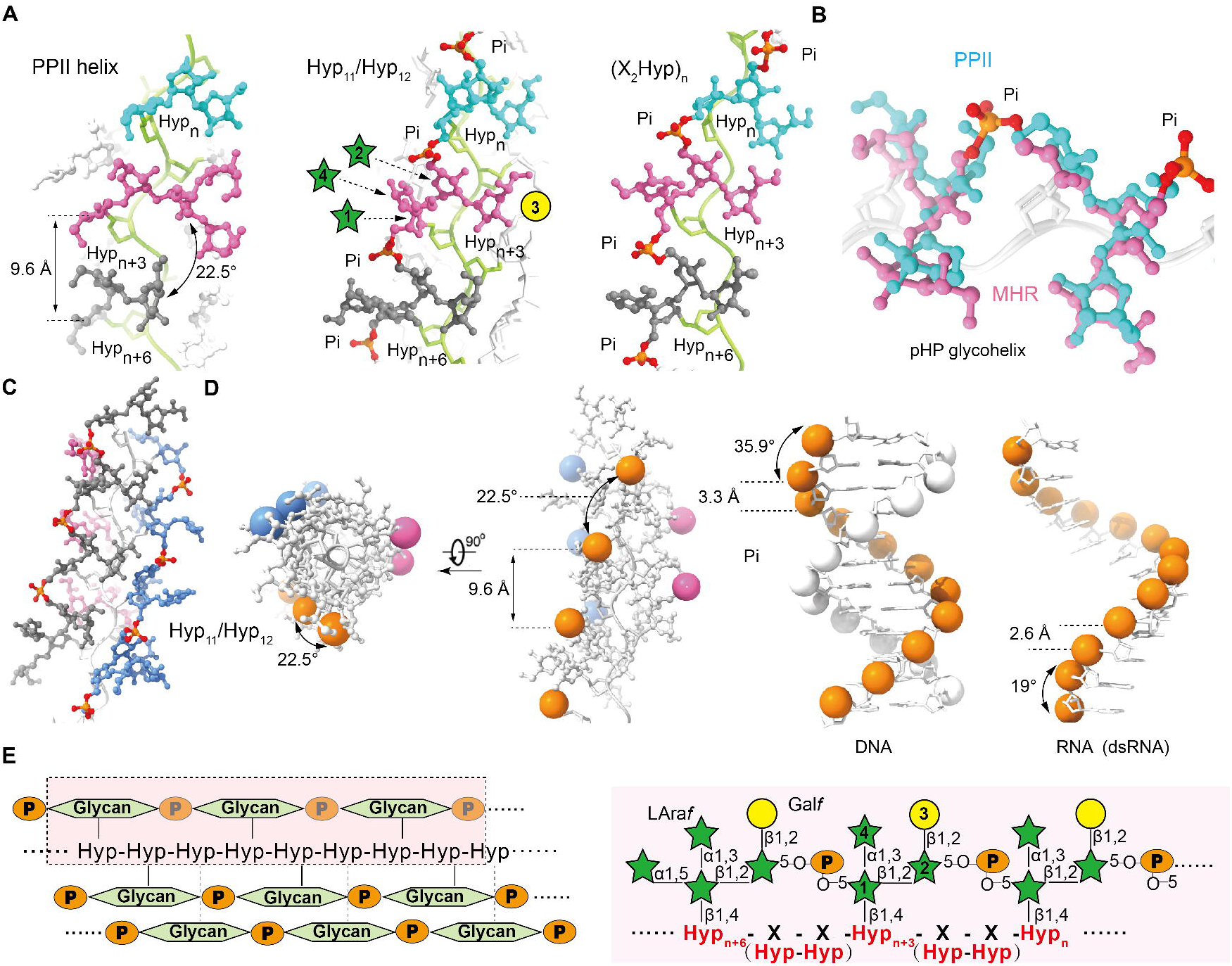
Folding paradigm of the Hyp-linked glycans in the mastigoneme. **(A)** Similar structural features of the arabinoglycans on the PPII helix of Mst1 and the pGlycans on Hyp_11_(X_2_Hyp)_n_ Hyp_12_ of Mstax-MHRs. The pGlycans attached to (X_2_Hyp)_n_ and Hyp_11/12_ share the same basic structure unit. In the (X_2_Hyp)_n_ segment, one array of the pGlycans lines one side of the polypeptide chain. In the Hyp_11/12_ segment, three arrays of the pGlycans, each of the same structure as shown on the left, form a shell surrounding the poly-Hyp chain. All Hyp-linked glycans share the same core composition of three Ara*f* and one Gal*f*. Glycan chains attached to Hyp residues n, n+3, and n+6 in structure are colored cyan, magenta and gray, respectively. Phosphorus atoms are colored orange. **(B)** Glycans attached to the PPII helix and pGlycans on Hyp_11/12_ share the same secondary structural element, which we name the poly-Hyp (pHP) glycohelix. Glycopeptides from the PPII helix (cyan) and Hyp_11/12_ (magenta) are superimposed. The pHP glycohyelix refers to the structure of the glycans attached to Hyp residues n and n + 3. **(C)** Three arrays of the pHP glycohelices constitute the high-order glycoshell surrounding the linear poly-Hyp peptide chain. Each pHP glycohelix is identified by the same color. Phosphorus atoms are colored orange. **(D)** Comparison of the 5’,5’-phosphodiester helical parameters in the pGlycan chain with the 3’,5’-phosphodiesters in the DNA and RNA double helix. The phosphate groups are highlighted as spheres. The ideal DNA and RNA helices were generated in AlphaFold3 (54). For visual clarity, only one strand of the RNA double helix is shown. **(E)** Schematic summary of the folding of pGlycans. *Left*: The general folding patterns of three pHP glycohelices of the pGlycans. *Right*: Common core sugar moieties in the pGlycans that fold to a pHP glycohelix. Ara*f*, Gal*f*, and phosphate are represented by green pentagons, yellow circles, and orange circles, respectively.

The PPII helix conventionally refers to the poly-Pro peptide, which is a protein secondary structure. In the mastigoneme, it is Hyp, instead of Pro, to which the glycans are attached. To distinguish the poly glycol-Hyp structure from the simpler PPII helix, we propose to name the secondary structure of the glycans attached to every three Hyp residues as the poly-Hyp (pHP) glycohelix. It is not surprising that the basic parameters for the pHP glycohelix are partly defined by the polypeptide to which the glycans are bound (Fig. 4A, B). Three arrays of the pHP glycohelices assemble to result in the high-order glycoshell structure surrounding the linear peptide chain of poly-Hyp (Fig. 4C).

## Discussion

To date, our understanding of the three-dimensional structures and folding principles of glycans remains disproportionally limited. In addition to the inherent complexity of monosaccharide diversity, branching patterns, stereo-specificity and regiochemistry, the chemical nature of glycosidic bonds adds another layer of complexity, making the folding of glycans more complex than that of proteins. Unlike peptide bonds, which form sp^2^-hybridized plane units that restrict conformational flexibility and lead to predictable structural folds, glycosidic bonds involve monosaccharides with sp^3^-hybridized carbon atoms and multiple hydroxyl groups. The sp^3^ hybridization of these hydroxyl oxygens further increases the conformational freedom of glycans.

The heavily glycosylated *Chlamydomonas* mastigoneme affords an excellent opportunity to probe high-order assembly of glycans. At the high resolution, we can now confidently describe the packing of inter- and intra-glycan chains, marking a significant milestone in unveiling the three-dimensional structures of glycans and glycoconjugates (Fig. 4).

The discovery of covalent bonds linking adjacent glycan branches highlights the complexity of glycan organization. These bonds, which have not been observed in native molecules before, are analogous to the 3’,5’-phosphodiester bonds that link nucleotides in DNA and RNA (Fig. 4D). In the glycosylated Hyp_11/12_ segments, the three strings of 5’,5’-phosphodiesters also appears in the form of a triple helix, though with a much larger helical rise compared to that in DNA and RNA (Fig. 4C-E).

Interestingly, the structural organization of arabinoglycosylated poly-Hyp appears to be independent of the phosphodiester bonds. In Mst1, glycans derived from Hyp residues n and n+3, whether linked by phosphodiester bonds or not, show similar packing patterns (Fig. 4A, B). This raises the question: what is the function of these bonds? The linkage between the two 5’-OH groups of neighboring arabinose residues, which belong to two different glycan chains, restricts the configuration of these hydroxyl groups. As a result, the 5’,5’-phosphodiester-linked structures likely enable more specific interactions with binding partners. Given that Mstax serves as the central shaft, a relatively more rigid sugar shell in the stem region may be important for transmitting motion along the 600-nm mastigoneme. Furthermore, the phosphate group increases local electronegativity and participates in the coordination of a cation at the interface between Mstax and Mst1 (Fig. 3B). As to the PPII helix of Mst1, the Ser-Hyp_3_ repeat occurs more frequently than other Ser-Hyp_3+_ repeats. These Ser-Hyp_3_ repeats may lose the ability to form the 5’,5’-phosphodiester bond by disrupting the local geometry required for the phosphodiester bond formation between adjacent repeats.

The broader distribution of 5’,5’-phosphodiester bonds and their physiological functions, beyond their structural role, remain to be investigated. More importantly, the enzyme responsible for catalyzing these bond formations has yet to be identified. The structure-based discovery of the unprecedented 5’,5’-phosphodiester bonds suggests that the complexity of glycan organization may exceed our imagination. Notwithstanding these puzzles, our attempt to define the secondary structural elements for glycans represents an important step towards a systematic investigation of the structures of glycans and glycoconjugates. The analysis reported here set the foundation for understanding glycan folding.

## Acknowledgements

We thank N. Zhou and J. Lei for technical support during EM data collection. We thank the Tsinghua University Branch of China National Center for Protein Sciences (Beijing) for providing the cryo-EM facility support and the computational facility support. We thank D. Yan and H. Deng in Center of Protein Analysis Technology, Tsinghua University, for MS analysis. This work was funded by the National Natural Science Foundation of China (32330052, N. Y, 32341016, 32171204, C.Y; 32370813, 31991191, J. P.), the National Key R&D Program of China (2020YFA0509301, C.Y., 2018YFA0902500, J. P.), Beijing Frontier Research Center for Biological Structure, Beijing Advanced Innovation Center for Structural Biology, State Key Laboratory of Membrane Biology, Tsinghua University Initiative Scientific Research Program, and Start-up funds from Tsinghua-Peking Center for Life Sciences and Tsinghua University.

## Author contributions

J.H., and H.T. prepared EM samples for data collection. H.T. prepared NMR samples for data collection. J.H. collected the cryo-EM data. J.H., and C.Y. processed the cryo-EM data and determined the structure. J.H., N.Y., and C.Y. did structural analysis. J.P., C.Y., J.H., and T.H. provided biochemical analysis. Y.N., J.P., and C.Y. provided resources. N.Y., J.H., and C.Y. wrote the paper with input from all authors.

